# Prefrontal, striatal, and VTA subnetwork dynamics during novelty and exploration

**DOI:** 10.1101/2021.11.24.469851

**Authors:** Adam J.O. Dede, Nader Marzban, Ashutosh Mishra, Robert Reichert, Paul M. Anderson, Michael X Cohen

## Abstract

Multiple distinct brain areas have been implicated in memory including the prefrontal cortex (PFC), striatum (STR), and ventral tegmental area (VTA). Information-exchange across these widespread networks requires flexible coordination at a fine time-scale. In the present study, we collected high-density recordings from the PFC, STR, and VTA of male rats during baseline, encoding, consolidation, and retrieval stages of memory formation. Novel sub-regional clustering analyses identified patterns of spatially restricted, temporally coherent, and frequency specific signals that were reproducible across days and were modulated by behavioral states. Clustering identified miniscule patches of neural tissue. Generalized eigen decomposition (GED) reduced each cluster to a single time series. Amplitude envelope correlation of the cluster time series was used to assess functional connectivity between clusters. Dense intra- and inter regional functional connectivity characterized the baseline period, with delta oscillations playing an outsized role. There was a dramatic pruning of network connectivity during encoding. Connectivity rebounded during consolidation, but connections in the theta band became stronger, and those in the delta band were weaker. Finally, during retrieval, connections were not as severely reduced as they had been during encoding, and specifically theta and higher-frequency connections were stronger. Underlying these connectivity changes, the anatomical extent of clusters observed in the gamma band in the PFC and in both the gamma and delta bands in the VTA changed markedly across behavioral conditions. These results demonstrate the brain’s ability to reorganize functionally at both the intra- and inter-regional levels during different stages of memory processing.

**SIGNIFICANCE STATEMENT:** The brain is often thought of as a mosaic of areas each with static functions that activate or deactivate with task demands. Here, we used large-scale recordings (196 simultaneous electrodes) and developed a multivariate analysis approach to analyze data from all our recording locations simultaneously. This analysis revealed that the brain dramatically reorganized itself at both local and long-distance spatial scales during different stages of memory processing. These results demonstrate an extreme degree of flexibility in functional anatomy. Rather than thinking about the brain as a set of static mosaic tiles, it may better be characterized as a quickly moldable piece of clay where each part’s function changes as the whole is reshaped from moment to moment.

## INTRODUCTION

The study of memory has long been guided by the goal of defining the functions of various brain regions (Squire and Dede, 2015; Lashley, 2020), yet it is increasingly recognized that the storage, retrieval, and active use of memory is supported by a distributed network including the prefrontal cortex (PFC), striatum/basal ganglia (STR), and hippocampus (HPC). For example, voxel-level analyses of fMRI data have revealed widely distributed semantic, episodic, and working memory representations (Huth et al., 2012; Rissman and Wagner, 2012). Simultaneous physiological recordings from multiple areas in monkeys engaged in memory tasks have indicated complex inter-regional coordination (Constantinidis and Procyk, 2004; Loonis et al., 2017), and optogenetic manipulations in rodents have demonstrated a clear interaction between the medial PFC and HPC for memory retrieval (Rajasethupathy et al., 2015). Beyond memory-specific studies, coordinated activity between brain regions is widely believed to allow neural circuits to flexibly bind cell assemblies and efficiently orchestrate information transfer (Singer, 2009; Jensen and Mazaheri, 2010; Wang, 2010).

The nature of interactions between regions varies as a function of task demands. In some cases, regions appear to cooperate (Turk-Browne et al., 2009; Wimmer and Shohamy, 2012), but in others, they appear to compete (Packard and McGaugh, 1996; Wimmer et al., 2014; Loonis et al., 2017). More generally, it is unclear how these interactions are mediated, and how interactions may be different at different times during memory formation and use.

Dopamine (DA) has been strongly implicated in synchronization and network-level dynamics (Montaron et al., 1982; Williams et al., 2002; Costa et al., 2006; Dejean et al., 2012). DA stems from the ventral tegmental area (VTA) and substantia nigra, and projects widely to most of the brain, with the densest projections into the STR, PFC, and HPC (Otmakhova et al., 2013; Kafkas and Montaldi, 2018; Kaminski et al., 2018). The VTA is therefore positioned to facilitate the coordinated processing that allows the brain to generate a memory-guided action plan from moment to moment (Fujisawa and Buzsaki, 2011; Jo et al., 2013; Beeler and Kisbye Dreyer, 2019; Freedberg et al., 2020).

Here, we utilized rats that had been implanted with high-density recording arrays in the STR, VTA, and PFC as part of a separate project studying reward-learning. We investigated how intra- and inter-regional dynamics varied as a function of behavioral state during a memory task. Rats were exposed to a simple novel-object memory paradigm, likely sensitive to lesions in the hippocampal system (Buffalo et al., 1999; Mumby, 2001; Aggleton and Brown, 2006). Given that previous research has associated theta frequency oscillations with influence from the hippocampus (Buzsáki and Draguhn, 2004; Buzsáki, 2006), we investigated whether network structure reflected increased influence from theta signaling during any phase of memory processing. More generally, we developed a mix of novel data-mining and hypothesis-driven network analyses to increase our understanding of the mechanisms of inter-regional connectivity.

## METHODS

Analysis was primarily carried out using custom written MATLAB code. ANOVA tests and some figure generation was carried out in R.

### Experimental Design

The experimental procedures have been described previously (Mishra et al., 2020). Briefly, all experimental procedures were performed in accordance with the EU directive on animal experimentation (2010/63/EU), and the Dutch nationally approved ethics project 2015-0129. All recordings were performed in the lab of MXC. We included five male Long-Evans TH:Cre rats (∼3 months old, weight: 350-450 g at time of recordings). Non-overlapping findings from this dataset have been reported elsewhere (Mishra et al., 2020).

Electrophysiological recordings were collected from the PFC, STR, and VTA. There were 64 contacts per region. For target recording locations see Figure 2a. 64 electrodes covered an area of 1 x 2 mm with typical spacing of 225 *μ*m in each shank and 330 μm between shanks in PFC. STR electrodes also covered an area of 1 x 2 mm with the same shank distance (330 μm). However, two shanks contained only tetrodes and two shanks had only single sites with typical spacing of 130 *μ*m between single sites and 660 *μ*m between tetrodes. VTA implants contained 8 shanks of 8 electrodes each and covered an area of 1.5 x 0.14 mm.

**Fig. 2.**
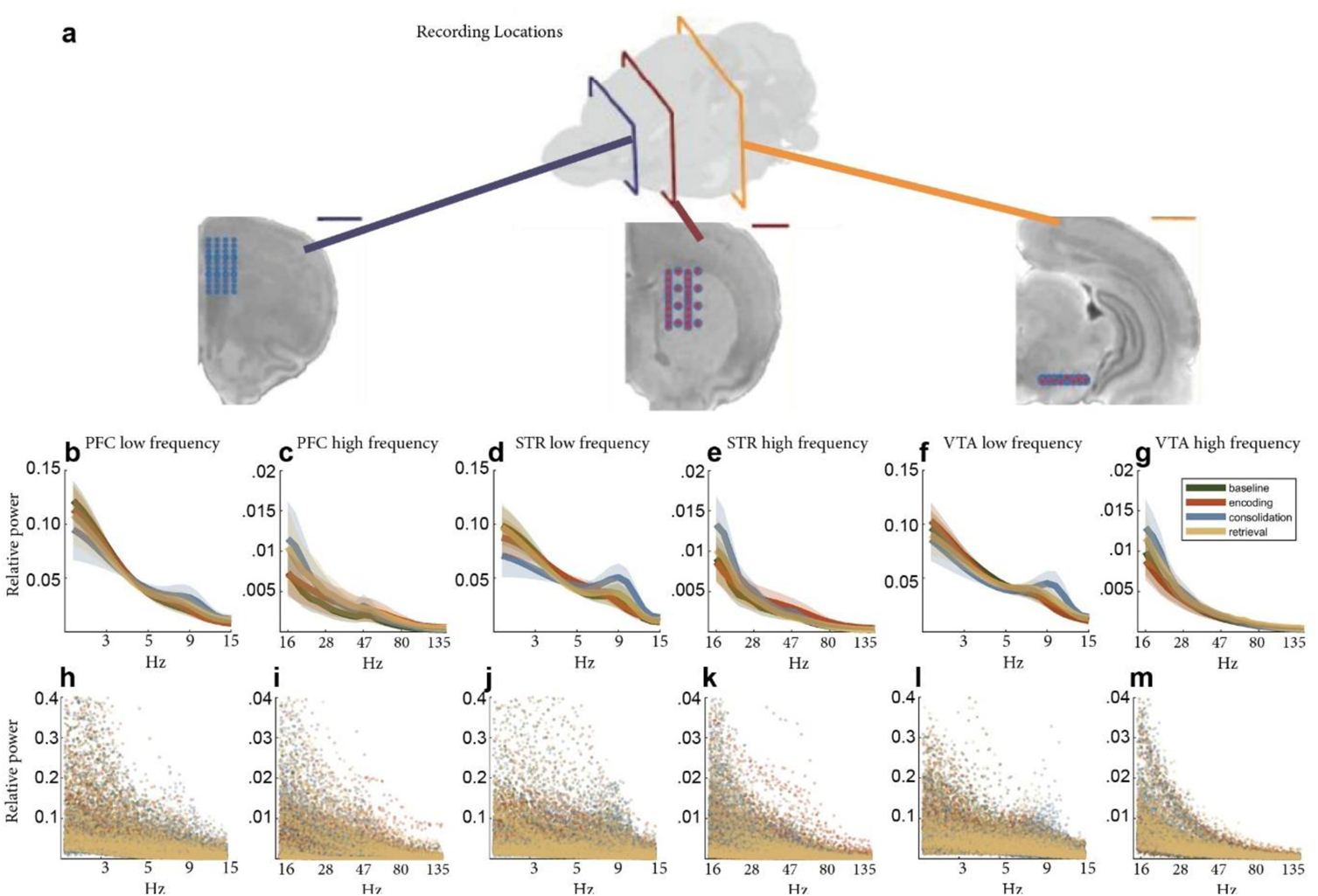
Power spectra do not differ reliably between conditions. **a** Recording locations are shown for the PFC (left), STR (middle), and VTA (Right). Scale bars indicate 2mm. **b-g** Group mean relative power spectra are displayed. Power spectra were calculated for each channel. Channel spectra were averaged and normalized to the summed spectral power across frequencies within each animal and region. Shaded regions indicate standard error of the group mean. h-m Relative power is shown for every channel individually, which highlights the variability in the spectra of individual channels.

After habituation, each experimental session consisted of four conditions. First, animals were placed in an open field. Second, a novel object (e.g., a cup or toy) was presented in the middle of the box. Third, the animal was alone in the open field again. Fourth, the same object presented in the second condition was presented again. Each condition lasted between five and six minutes. We termed these conditions, baseline, encoding, consolidation, and retrieval, respectively. Rats moved freely throughout experimental sessions. There was no delay between conditions. A camera was placed above the box to track movement (Figure 1a-c). A maximum of one session per animal was recorded on a single day. There were 28 recording sessions in total.

**Fig. 1.**
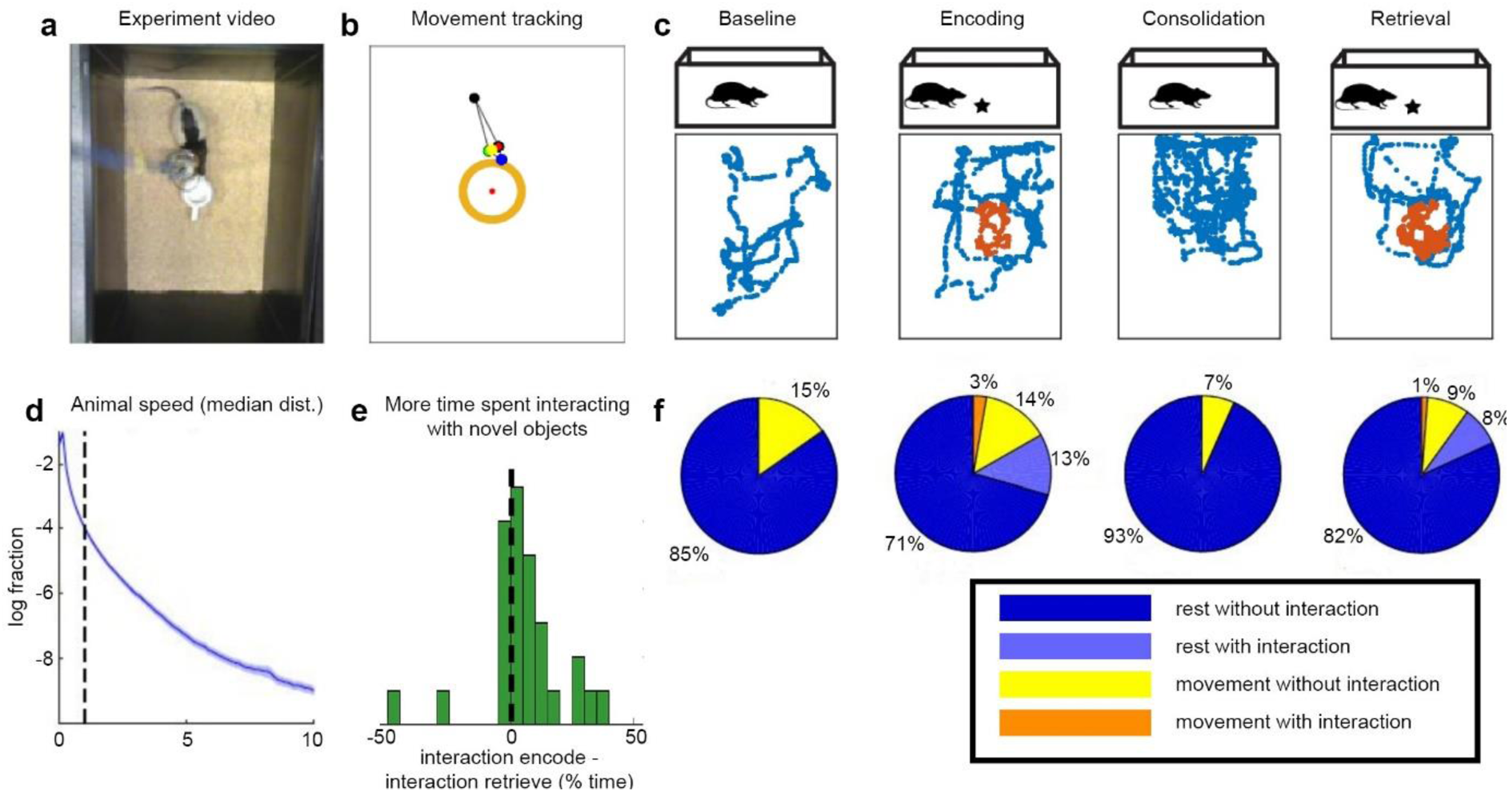
Behavioral paradigm and behavior results. **a** Still image taken from video recording of an experimental session. The rat is exploring a white object. **b** Output of movement tracking results for the frame shown in panel **a**. **c** Example experimental session behavioral data. Stars indicate the presence of an object to explore (encoding and retrieval periods). During baseline and consolidation periods, there were no objects in the box. Different objects were used on different testing days. Within day, the same object was used in the encoding and retrieval periods. In the bottom of each panel is the path followed by the rat during the corresponding condition. Orange versus blue points differentiate locations with and without interaction with the object, respectively. **d** Median distribution of animal speed movement from all the recordings. The dashed line shows the motion speed threshold separating resting from movement. **e** Histogram depicts memory for the object in terms of the percentage of time spent interacting with the object during the encoding period minus the corresponding percentage during the retrieval period. In general, more time was spent interacting with the object when it was novel (after removal of outliers more than 2 SDs below mean *t*(25)=4; *p*<<.001). **f** Pie charts show percentages of time spent in different behavioral states during each of the behavioral conditions.

Using data from video recordings and DeepLabCut (Mathis et al., 2018), we created binary vectors indicating interaction with the object (during encoding and retrieval conditions) and movement. These were upsampled to 1000 Hz and aligned to LFP data.

### Statistical Analyses: Calculating memory strength

For each session, the percentage of time spent interacting with the object was calculated for the encoding and retrieval periods. The percentage during retrieval was subtracted from the percentage during encoding. Positive values indicate that the rat spent more time exploring the object when it was novel.

### Local field potential data cleaning

Data were notch filtered to remove 50 Hz line noise, ICA filtering was done and components that appeared to capture muscle and line noise were removed, channels that appeared to be contaminated with noise by visual inspection were removed. Finally, cross-channel covariance matrices were calculated in 2000 ms windows in steps of 100 ms. A mean covariance matrix was calculated. Epochs whose covariance matrices were more than 2 standard deviations from the mean were discarded from further analysis. Distance between epoch and mean covariance matrices was measured using matrix Euclidean distance.

### Identifying intra-regional clusters

Data were filtered using 42 logarithmically spaced central frequencies between 2 and 150 Hz. After filtering, data from novel and repeat object periods were limited to periods of interaction with the object. Electrode X electrode correlation matrices were calculated in non-overlapping 2.5s epochs. These epochs were averaged together to create a single electrode X electrode correlation matrix (Figure 3a). In addition, the average correlation matrix was calculated 20 additional times with an evenly-spaced sliding window of 10% of the data left out from each average. These 20 partial averages were used for validation. This was done for each behavioral condition independently.

**Fig. 3.**
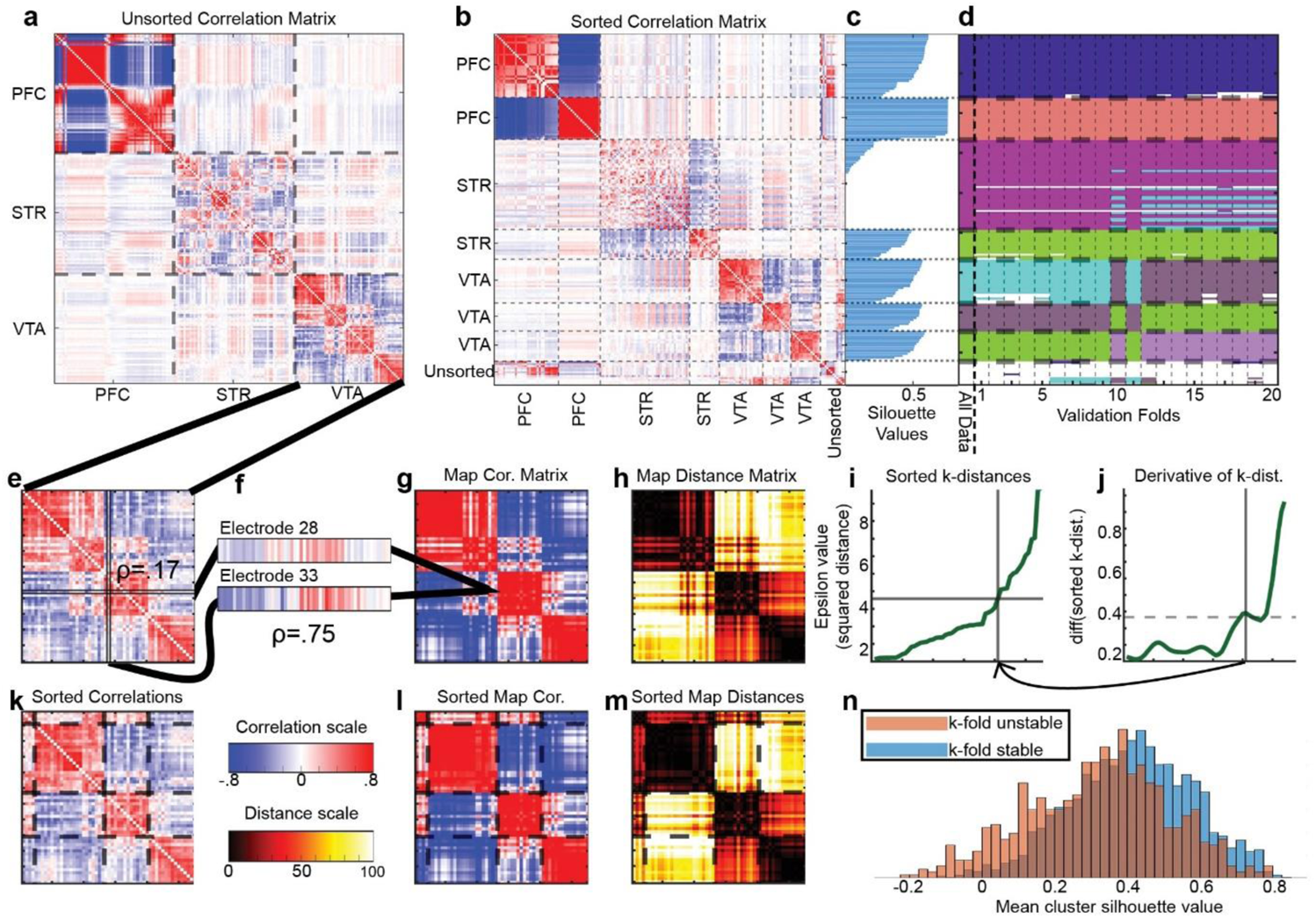
Clustering methods and validation. **a** Unsorted channel X channel correlation matrix for rat 5 during the baseline period constructed using data narrowband filtered at 5.2 Hz. **b** The same set of correlations after application of sorting pipeline. **c** Silhouette values associated with each channel. One cluster in the STR had low silhouette values, indicating poor clustering (this cluster was removed from subsequent analyses). **d** Clusters detected in “All Data” (far left column) and in each of 20 validation folds. Different pseudo-colors indicate different clusters. The channels comprising the cluster with low silhouette values are not always clustered together, indicating instability (pink with blue stripes) (note that an entire cluster can switch colors in different folds; the important metric is whether the color is homogeneous across channels within the cluster). **e-m** These panels display the clustering pipeline. **e** Each region’s correlation matrix is considered separately. The VTA is shown here. **f** The correlation between each row and column of the correlation matrix is calculated. Channels from electrodes 28 and 33 are displayed as examples. **g** These correlations are organized in a matrix that encodes similarities of connectivity profiles, rather than bivariate correlations. **h** This connectivity-profile correlation matrix was transformed into a Euclidean distance matrix to increase contrast and enforce positivity. **i** The k-distance for each channel represents how far (epsilon; y-axis) one would have to go in units of squared distance (panel **h**) in order to find k nearest neighbors. K is set to 8. Values are sorted from smallest to largest. **j** The derivative of the k-distances was approximated by taking the running difference between pairs of k-distances. The horizontal dashed line indicates the detection threshold for sharp discontinuity. The vertical solid line indicates a local peak. The arrow pointing back to panel **i** shows how the detected index in the derivative is used to select the corresponding epsilon value. This epsilon is used as input to the DBscan algorithm for clustering. **k-m** Correspond to the matrices shown in panels **e**,**g**, and **h**, but with channels sorted according to the result of the DBscan clustering. Cluster borders are indicated with dashed lines. **n** Mean silhouette values across channels in clusters that were stable across 20-fold validation (blue) and in clusters that were not stable across 20-fold validation (red). Stability was not determined using silhouette value (see methods), and there was no mathematical necessity that stable clusters would be expected to have higher silhouette values.

Clustering was done for each animal, condition, region, frequency, and validation fold independently. Before clustering, we first took the correlation coefficient of each row of the channel X channel matrix compared to each column of the matrix (Figure 3e-g). The resulting matrix was the same size as the input matrix, but now values in the matrix represented how the map of connectivity associated with one channel correlated with the map of connectivity associated with another channel (Liu et al., 2012). Finally, we took the squared Euclidean distance comparing each row to each column of the new matrix (Figure 3h). Squaring accentuates high similarities and forces all values to be positive, both of which facilitate clustering. This final matrix is referred to as the distance matrix.

The DBscan algorithm (Ester et al., 1996) was applied to distance matrices. The DBscan algorithm requires two input parameters: K and epsilon. Epsilon is the search radius around each point. K is the number of points that must be found within that radius in order for a given point to be considered a central point in a cluster. We chose the value of k to be constant at 8 for all clustering. This was done for two reasons. First, Ester et al. (Ester et al., 1996) noted that cluster discovery is largely invariant to the choice of K within a reasonable range. Second, we tested all values of K between 2 and 22 and visual inspection of resulting silhouette values of clustering schemes suggested that k=8 was reasonable (Extended data 2-1).

The silhouette value is a measure both of how well a point fits into a particular cluster and how poorly it fits into any other cluster. A good cluster organization will yield clusters that maximize the fit of all points to their respective clusters while minimizing the fits of points to other clusters (Rousseeuw, 1987; Tan et al., 2018).

Our choice for the epsilon value was set dynamically for each run of DBscan. To do this, the 8-distance values were calculated. 8-distance refers to the minimum epsilon value needed in order to reach 8 points from a given point to be clustered. When all 8-distances in a data set are sorted and these values are plotted, natural divisions in the cluster structure of the data can be identified at points of sharp steepness in the 8-distance plot (Figure 3i). Algorithmic identification of sharp steepness was identified using the running difference between sorted 8-distance values (Figure 3j). The running difference between sorted 8-distance values approximates the first derivative of the curve, so peaks in the plot correspond to points of maximum steepness in the 8-distance values. We identified the first peak above a threshold for each clustering run. The threshold was the mean of the running difference plus 2 standard deviations. Threshold calculation excluded the maximum value and the surrounding 5 points on either side. The epsilon value corresponding to the detected peak was used for clustering (see vertical and horizontal lines in panels i and j of Figure 3).

For each animal, condition, region, and frequency, clustering was performed on the correlation matrix calculated from the full recording and also on each of the 20 validation folds. Each cluster was examined across folds individually. For each fold, we asked what proportion of the channels in the cluster in the full data set were clustered together in the fold. We termed this value the agreement value. We further asked what proportion of the channels that were not a part of the cluster in the full data set were also given the same label as that which yielded the highest agreement value. We termed this value the outside value. The agreement value minus the outside value was termed the net agreement value, and clusters with an average net agreement value below .85 across folds were discarded as unstable (Figure 3d).

### Aggregating clusters

Normalized mutual information (NMI) was calculated for all pairs of cluster schemes within each region using equation 3 from Strehl and Ghosh (Strehl and Ghosh, 2002). NMI yields a measure of the similarity between two cluster schemes of the same data. It is robust to differences in arbitrary labels (e.g. cyan vs. mauve in Figure 5d) and to missing data (e.g. unclustered white electrodes in Figure 5d). NMI ranges from 0 to 1. Values near 0 represent completely different clustering schemes where channels grouped into the same cluster in one scheme are in different clusters in another scheme. An NMI of 1 indicates an identical cluster scheme. NMI was calculated between cluster schemes from within the same condition. NMI values were averaged across conditions within each region, yielding a frequency X frequency matrix of cluster similarity for each region. Based on visual inspection of these matrices (Figure 5a-c), we decided to break frequency up into 5 bands. The breakpoints for these bands were chosen by a greedy search algorithm. The algorithm began with 4 breakpoints spaced evenly across logarithmic frequency space. For each breakpoint, the average NMI within all frequency bands and between all frequency bands was calculated. The between-NMI was subtracted from the within-NMI. This net NMI value was calculated for all possible positions of the current breakpoint such that it was at least 3 frequencies away from the two breakpoints (or ends) on either side of it. The breakpoint was moved to the position with the maximum net NMI value. This loop was repeated until no breakpoint moves were made. While increasing the number of breakpoints from 3 to 4 markedly increased the final net NMI, only a marginal increase was found by increasing to 5, confirming the use of 4 breakpoints to create 5 frequency bands.

**Fig. 5.**
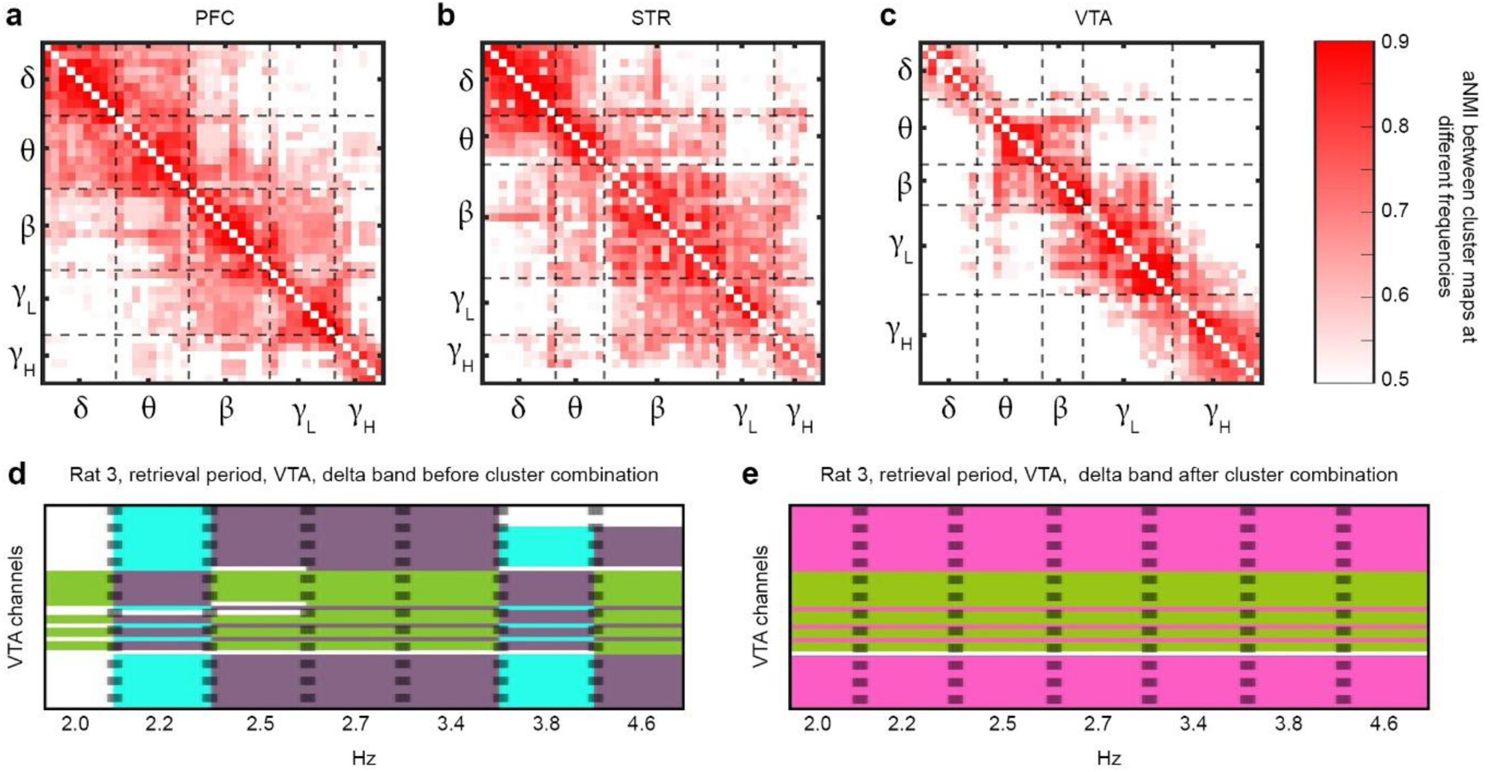
Defining frequency bands and combining clusters within bands. **a-c** Heat maps display the average normalized mutual information (aNMI) between cluster maps at different frequency bands. Averaging was done across rats and conditions. Dashed lines indicate the output of a greedy search algorithm that divided frequency space into bands. **d-e** Channel-by-frequency maps illustrating the aNMI-maximizing clustering results. The y-axis represents channel. The x-axis represents frequency. Pseudo colors indicate cluster groups. **e** A single cluster map has been constructed for all frequencies within the band such that aNMI between the final cluster map and the maps associated with the different frequencies within the band (panel **d**) has been maximized.

Next, cluster schemes were aligned within each frequency band for each rat, condition, and region independently. We used equation 5 from Strehl and Ghosh (Strehl and Ghosh, 2002) to calculate the average NMI (aNMI) between a candidate cluster scheme and all cluster schemes within a frequency band. The initial candidate cluster scheme was chosen by selecting the input cluster scheme that had the highest aNMI with the other cluster schemes within its frequency band. The initial candidate scheme was relabeled to meet two constraints: (i) λ_1_= 1; (ii) for all i =1, …, n − 1: λ_i_+1 ≤ max_j=1, …, i_ (λ_j_) + 1 (Strehl and Ghosh, 2002). Here, λ represents the cluster label of the electrode indicated by the subscript. Next, the algorithm looped over electrodes. For each electrode, the aNMI of the whole scheme was calculated with the electrode in question having each of the possible cluster labels available in the scheme. If the aNMI was higher for some other label than the electrode had at the start of the loop, then the electrode’s label was changed. Looping continued until no further changes were made. This yielded a single cluster scheme across the entire frequency band.

### Measuring changes in within-region functional structure

aNMI was used to measure cluster similarity between conditions (within frequency) and between frequencies (within conditions). In both ways of doing the analysis, each rat was considered independently. In the between-conditions analysis, cluster schemes from all four conditions were considered for one region and one frequency at a time. For each of these four cluster schemes (one from each behavioral condition), the aNMI was calculated with respect to the other three conditions. On this metric, values near 0 would indicate that within a particular frequency band, the functional structure of a region observed during a particular condition was dramatically different from other conditions. By contrast, values near 1 would indicate a high degree of functional stability between conditions. The values obtained from individual rats were subjected to a within-subjects ANOVA with the factors frequency band and condition. For conditions, dummy variables encoding linear contrasts were used to compare baseline vs. consolidation, encoding vs. retrieval, and periods with objects (encoding and retrieval) vs. periods without objects (baseline and consolidation). For frequencies, linear contrasts were used to compare delta vs. others, theta vs. others, low gamma vs. others, and high gamma vs. others.

The between frequency analysis was similar. Cluster schemes from all five frequencies were considered for one region and one condition at a time. For each of these five cluster schemes (one from each frequency band), the aNMI was calculated with respect to the other four conditions. Again, within-subjects ANOVA was used with the factors frequency band and condition. The same set of linear contrasts were used.

### Measuring changes in between-region functional connections

To facilitate measuring connections between regions, we used generalized eigendecomposition (GED) to reduce the signals from electrodes within each cluster to a single time series. The goal of GED is to identify a component, defined as a weighted combination of the channel time series from within each cluster, that maximizes the power of narrowband activity to broadband activity:

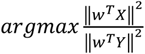

Where **X** is the narrowband-filtered data, **Y** is the broadband data, and **w** is the vector of channel weights. The solution to this optimization can be obtained from the GED on two covariance matrices: **S**=**XX^T^** and **R**=**YY^T^** (Parra et al., 2005; Cohen, 2021):

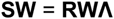

**W** is the square matrix of eigenvectors, and **Λ** is the diagonal matrix of eigenvalues. After solving the GED for each cluster, the eigenvector associated with the largest eigenvalue was used to calculate a weighted combination of the narrowband signals from the cluster resulting in a single time series for each cluster that explained the maximum amount of variance between the electrodes. The largest eigenvalue was divided by the sum of all eigenvalues in order to estimate the proportion of variance explained by the single time series (Figure 4f).

**Fig. 4.**
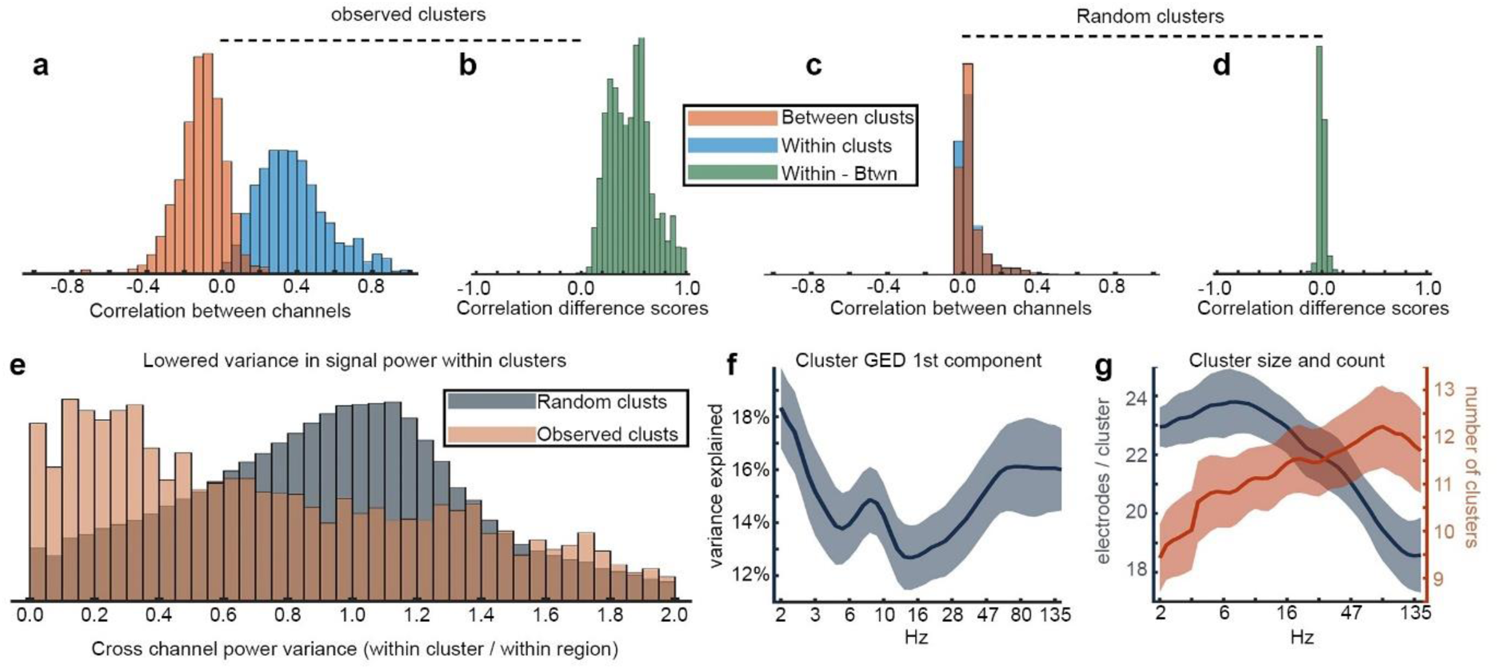
Characteristics of clusters. **a** Histograms show distributions of correlations for pairs of channels that were within the same cluster (blue) or between channels from different clusters (red). **b** The histogram shows the distribution of difference scores calculated by subtracting between-cluster correlations from within-cluster correlations. Subtractions carried out for correlation values from the same animal, region, condition, and frequency. **c-d** Similar to a and b, but clusters were chosen randomly. **e** Histograms show the distributions of variance in power between channels within a cluster divided by the variance in power between channels within the corresponding region. Lower values indicate that there is less variance in power within a cluster than would be expected given the variance in its containing region. The red histogram shows the values calculated for observed clusters. The grey histogram shows the values calculated for random clusters. **f** The first component of a generalized eigendecomposition (GED) performed on the channels within each cluster generally explained between 13% and 17% of between channel signal variance. The y-axis displays variance explained by the first GED component. The x-axis displays frequency. Higher values indicate that the entire cluster is well-characterized by a single time-series. **g** There was a larger number of smaller clusters detected at higher frequencies. The left y-axis displays the number of channels grouped into each cluster. The right y-axis displays the total number of clusters detected. The x-axis displays frequency. For panels **f** and **g**, shaded regions indicate standard error of the group mean.

Connectivity between cluster time series was assessed using amplitude envelope correlations (Bruns et al., 2000). Time series were transformed into amplitude envelopes by taking the absolute value of the Hilbert transform. For every pair of cluster time series within a given rat and condition, correlations were calculated in non-overlapping 2.5 second windows. Windows with a correlation greater than 2 standard deviations from the mean were ignored. The remaining correlations were averaged together to obtain a connectivity strength for the pair of clusters. To assess the significance of these connectivity strength values, the same amplitude envelope correlation analysis was carried out again with one of the time series offset such that the last X data points in the time series were cut from the end and placed at the beginning of the series where X was a random value. This recalculation was carried out 1000 times for each pair of connections. Connections whose original correlation was stronger than 950 or more of the comparison correlations were deemed significant.

For graph-theoretic measurements, each cluster was treated as a node and significant connections were treated as weighted edges. Strength, betweenness centrality, clustering coefficient, and average path length were calculated using functions from the Brain Connectivity Toolbox (Rubinov and Sporns, 2010). These measures were combined across rats within each condition and sorted by strength (Figure 7l-o). The total strength within each combination of frequency band and region was summed and plotted as a heatmap for each condition (Figure 7p-s). Summed strength values were submitted to a series of within-subject ANOVAs with frequency band, region, and condition as factors. ANOVAs compared two conditions at a time: baseline vs. encoding, baseline vs. consolidation, baseline vs. retrieval. t-tests were used to assess changes in total strength in the delta and theta frequency bands.

**Fig. 7.**
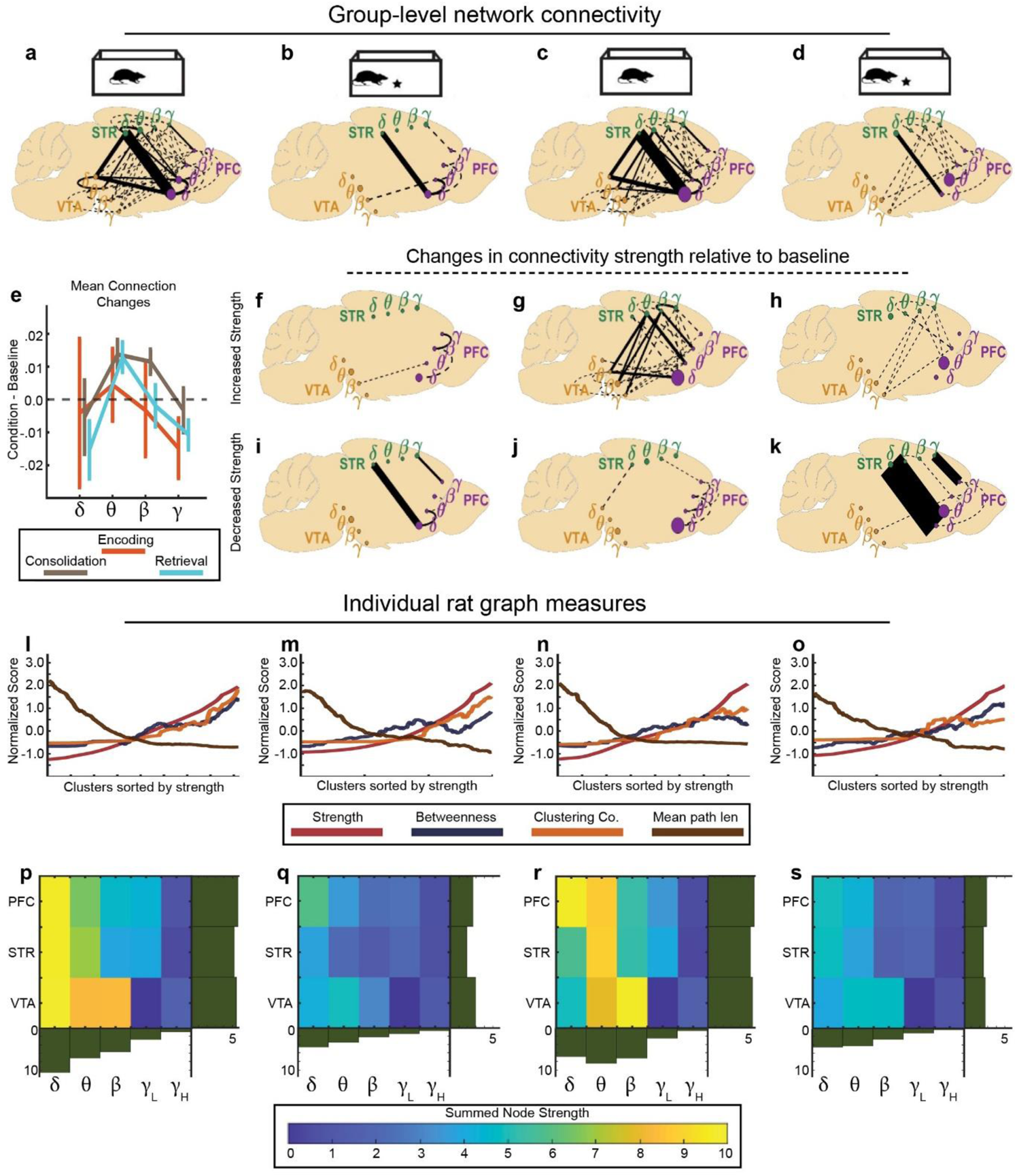
Dynamics of inter-regional connectivity across behavioral epochs. **a-d** Group mean correlations between signals derived from frequency-specific regional clusters are shown as line thickness. Solid lines range from ρ=.05 to ρ=.20. Dashed lines represent weaker connections (ρ>0.0). All visualized connections were significant at the individual level for at least 4/5 animals. Each animal contributed only its strongest single connection to each graph edge. Node size represents betweenness centrality. These connection strengths are reused in panels **e-k**. **e** Group mean change relative to baseline in connection strength for connections between nodes within different frequency bands. Inter-regional connections in the theta band had marginally increased strength in the consolidation and retrieval periods (t-test against 0, ps<.07). Connections in the beta band had increased strength in the consolidation period (p=.0503). **f-h** Specific connections that exhibited increased strength relative to baseline in the encoding (**f**), consolidation (**g**), and retrieval (**h**) periods. **i-k** Similar to **f-h**, but for decreased strength connections. Throughout f-k, solid lines represent connectivity changes of between ρ=.0125 and ρ=.07. Dashed lines represent weaker connections (ρ>0.0). **l-o** Graph theoretic measurements of each animal’s connectivity matrix were calculated for all significant connections (rather than taking only each animal’s strongest edge between any two nodes). Metrics were z-scored for display on a single scale. Node-metrics from all animals were combined and sorted by strength for plotting. In general, nodes with high strength also had high betweenness centrality, high clustering coefficients, and low mean path lengths. **p-s** Summed strength values of all nodes in different regions (y-axis) and within different frequency bands (x-axis). Marginal histograms display the mean value of their respective rows or columns of the heatmap. Overall node strength was lower in the encoding and retrieval periods (panels **q** and **s**), and the frequency of peak nodal strength shifted from delta to theta when comparing baseline (panel **p**) to consolidation (panel **r**).

To visualize connectivity maps, connections were pooled across animals. First, we took each rat’s strongest significant connection between pairs of regions and frequency bands. Because of a limited number of significant connections involving high gamma, low and high gamma were combined for this analysis. Next, for connections that were significant for at least 4/5 animals, the median connectivity strength across rats was calculated. This resulted in a group connectivity matrix that was 12 X 12 (3 regions X 4 frequency bands). For display, these connections were plotted on a schematic of the rat brain using line thickness to indicate connectivity strength (Figure 7a-d). In addition, the group 12 X 12 connectivity matrix for the baseline period was used as a reference, and plots were generated to display the subset of connections that increased in strength relative to baseline (Figure 7f-h) and decreased relative to baseline (Figure 7i-k). Finally, the mean connectivity strength relative to baseline was also calculated between nodes within each frequency band (Figure 7e), and these relative changes in connectivity strength were compared to 0 using t-tests.

We examined the relationship between connectivity strength and memory strength. To do this, amplitude envelope correlations were calculated on an individual session basis for connections that were significant in the group (significant at the individual level for 4/5 rats). The mean of these session-wise connectivity values was taken for each animal. In addition, each animal’s mean memory strength was calculated by taking the mean of its individual session memory strengths. Sessions with memory strength more than 2 standard deviations from the mean were discarded from this analysis (2/28 recording sessions). Correlations were calculated between mean memory strengths and mean connectivity strengths. This analysis yielded similar results when sessions were kept separate and correlations were calculated across all 26 sessions (after removal of 2 outlier sessions).

Finally, we repeated the generation of pooled connectivity maps treating each region as a single large cluster. The same band divisions that were used in the main clustering analysis were used here. The goal of this analysis was to see whether similar connectivity patterns would be discovered if the clustering process was skipped.

### Data availability statement

The data that support the findings of this study are available from the corresponding author upon reasonable request.

### Code accessibility

Key custom functions are available on Github (https://github.com/adede1988/subNetworkDynamics.git). Full processing and analysis code is available from the corresponding author upon reasonable request.

## RESULTS

### Behavior

Animals were serially exposed to (1) an empty open field, (2) the same open field with a novel object, (3) the empty open field again, and finally (4) the open field with the same object. These conditions were termed baseline, encoding, consolidation, and retrieval, respectively (Figure 1c). During the baseline and consolidation periods, rats tended to sit still (85% and 93% of the time, respectively; Figure 1f). During the encoding and retrieval periods, rats rested for somewhat less time (84% and 90% of the time, respectively). Rats spent more time interacting with the object in the encoding than retrieval period (16% vs. 9%, respectively), and subtracting the percent of time spent interacting with the object during the retrieval period from the corresponding percentage during the encoding period yielded a significant difference (after removal of outliers more than 2 SDs below mean t(25)=4; p<.001; Figure 1e).

### Local Field Potential Power Effects

We calculated power spectra averaged across time and electrode for each behavioral condition and each brain region (Figure 2b-g). Repeated measures t-tests were used to compare each condition to baseline for each frequency individually. In general, spectral dynamics in all three regions were characterized by a 1/f-like decrease in power with increasing frequency, and a peak in the theta range (5-10 Hz). The only reliable difference in the spectral profiles between behavioral conditions was a relative increase in power around 4 Hz in the STR during the encoding phase.

Closer inspection of the individual power spectra per electrode revealed considerable inter-electrode variability (Figure 2h-m). This suggests that the multielectrode arrays may have spanned multiple functionally distinct neural networks. We therefore proceeded to identify clusters of electrodes based on inter-electrode correlation matrices.

### Identification of intra-regional clusters

We identified clusters of channels based on similar patterns of inter-channel correlations of their LFP time series, which were identified using the DBscan algorithm.(Ester et al., 1996) The clustering method was applied separately per animal, brain region, task condition, and narrowband frequency between 2 and 150 Hz, and the robustness of clusters was confirmed using 20-fold cross-validation (see Methods for details and Figure 3). Correlation matrices had strong block-diagonal patterns both between- and within-region, and these patterns were successfully detected and emphasized using clustering analysis (Figure 3a-b). Most clusters exhibited high silhouette values (Rousseeuw, 1987)(Figure 2n). Across animals, there was a similar number of clusters detected for each condition (range 270-283 clusters over all frequencies), and the number of clusters per condition did not vary widely between animals (range 250-285). However, not all clusters survived 20-fold validity testing.

### Cluster Validity and descriptive statistics

To ensure cluster validity, we assessed clusters in 20 validation folds. For each fold, we repeated the clustering analysis using only 90% of the data. Clusters that were not at least 85% consistent across folds were discarded as unstable (see methods). This procedure led to the elimination of 13.8% of clusters. Although we did not explicitly use silhouette values as a criterion for thresholding, eliminated clusters had lower average silhouette values than accepted clusters (Figure 3d and n). After validation, there was still no marked difference in clusters per condition (range 229-247), but rat 1 exhibited fewer stable clusters than other animals (rat 1: 199; range excluding rat 1: 238-261). The group average number of clusters summed over conditions and frequencies was not markedly different across regions (PFC: 77; STR: 76; VTA: 85.6).

Considering the narrowband signals used for clustering, the Pearson *ρ* values comparing channels within the same cluster were higher than those obtained when comparing channels from different clusters or that were unclustered (t-test on animal means: t(8)=10.7; p<.001; Figure 4a). The mean *ρ* value within clusters was .38, and the mean value between clusters was -.11. For 97% of cases, the average within cluster correlation was larger than the average between cluster correlation (Figure 4b). For each animal X condition X region X frequency, 1000 random cluster schemes were chosen with the same number of clusters and the same number of channels per cluster as those detected in our main analysis. Randomly chosen clusters did not exhibit a difference for within versus between cluster channel time-series correlations (Figures 4c-d).

The high correlation between channels within clusters suggested that clustering successfully detected groups of electrodes influenced by the same signal. To explore this further, we calculated the variance in power between channels within each cluster divided by the variance in power between all channels within each cluster’s region (Figure 4e). For random samples from a normal distribution, variance is insensitive to sample size, so this ratio would be expected to equal 1. Indeed, for randomly chosen clusters with the same frequency, region, and channel count characteristics as those observed, the average value for this ratio was 0.99. However, for observed clusters, the average value for this ratio was 0.83. The distributions of these power variance ratios were different (Two-sample Kolmogorov-Smirnov test: D=.24; p<<.001). We also found that a sizable percent of the variance between channels within each cluster could be explained by a single generalized eigendecomposition (GED) component (group average between 13% and 17% across frequencies; Figure 4f). Interestingly, there was a visually apparent local maximum in variance explained by the first GED component in the theta range (6-10 Hz).

Finally, the number of channels in any given cluster tended to be lower at higher frequencies, and correspondingly the number of clusters detected tended to be higher at higher frequencies (Figure 4g; ρ=-.72; p<<.001). This pattern suggests that the anatomical organization of higher frequency signals is more locally differentiated than that of low frequency signals.

### Aggregating clusters

In total, this procedure yielded a mean of 238.4 statistically reliable clusters per condition across animals. Visual inspection of electrode groupings revealed that clusters were largely stable across wide ranges of frequencies (e.g. Figure 5d). To assess this stability quantitatively, we calculated the normalized mutual information (NMI)(Strehl and Ghosh, 2002) between cluster schemes at different frequencies and averaged the resulting NMI matrix across conditions and animals for each region (Figure 5A-C). Based on NMI, we utilized a greedy optimization algorithm to select divisions between frequency bands that maximized average NMI (aNMI) within bands and minimized aNMI between bands. We divided the frequency space into 5 bands for each region (dashed lines in Figure 5A-C; see methods). Remarkably, despite the algorithm being applied separately per region and without a priori constraints regarding the size or spectral extent of clusters, the resulting frequency bands were similar across regions and corresponded to canonical frequency bands. In the PFC and STR the clusters mapped onto canonical delta, theta, beta, low gamma and high gamma (Figure 5A-B). By contrast, in the VTA there was a separate band for alpha, and beta was combined with low gamma (Figure 5C). For ease of explanation, the same band labels will be used throughout the text (Table 1).

**Table 1.** Divisions between frequency bands, values in Hz

| Region | Delta ($\delta$) | Theta ($\theta$) | Beta ($\beta$) | Gamma low (γL) | Gamma high (γH) |
| --- | --- | --- | --- | --- | --- |
| PFC | 2-4.6 | 4.6-12.0 | 12.0-34.3 | 34.3-79.4 | 79.4-150 |
| STR | 2-4.6 | 4.6-8.7 | 8.7-38.2 | 38.2-79.4 | 79.4-150 |
| VTA | 2-3.8 | 3.8-8.8 | 8.8-14.8 | 14.8-47.1 | 47.1-150 |

Next, information from different cluster schemes within each band was used to create a single cluster scheme within each band for each animal, condition, and region. To do this, we again used a greedy optimization algorithm. This time, the algorithm selected a cluster scheme that maximized the aNMI calculated across the cluster schemes within each band (Figure 5d-e).(Strehl and Ghosh, 2002) This procedure resulted in an average of 31.5 clusters per animal in each condition.

### Changes in within-region functional structure

Clusters were detected independently within-frequency and within-condition, and the steep drop-off in aNMI values away from the diagonals in Figures 5a-c indicates that cluster schemes were different in different frequency bands. To quantify cluster organization similarity across behavioral conditions and frequencies, we calculated the aNMI between pairs of cluster schemes detected either within a single frequency band but between different behavioral conditions (Figure 6a-f), or within a single condition but between different frequency bands (Figure 6g-m). An aNMI near 1 indicates that network structure is very stable across either frequency or condition, and an aNMI near 0 indicates that network structure is very different across either frequency or condition.

**Fig. 6.**
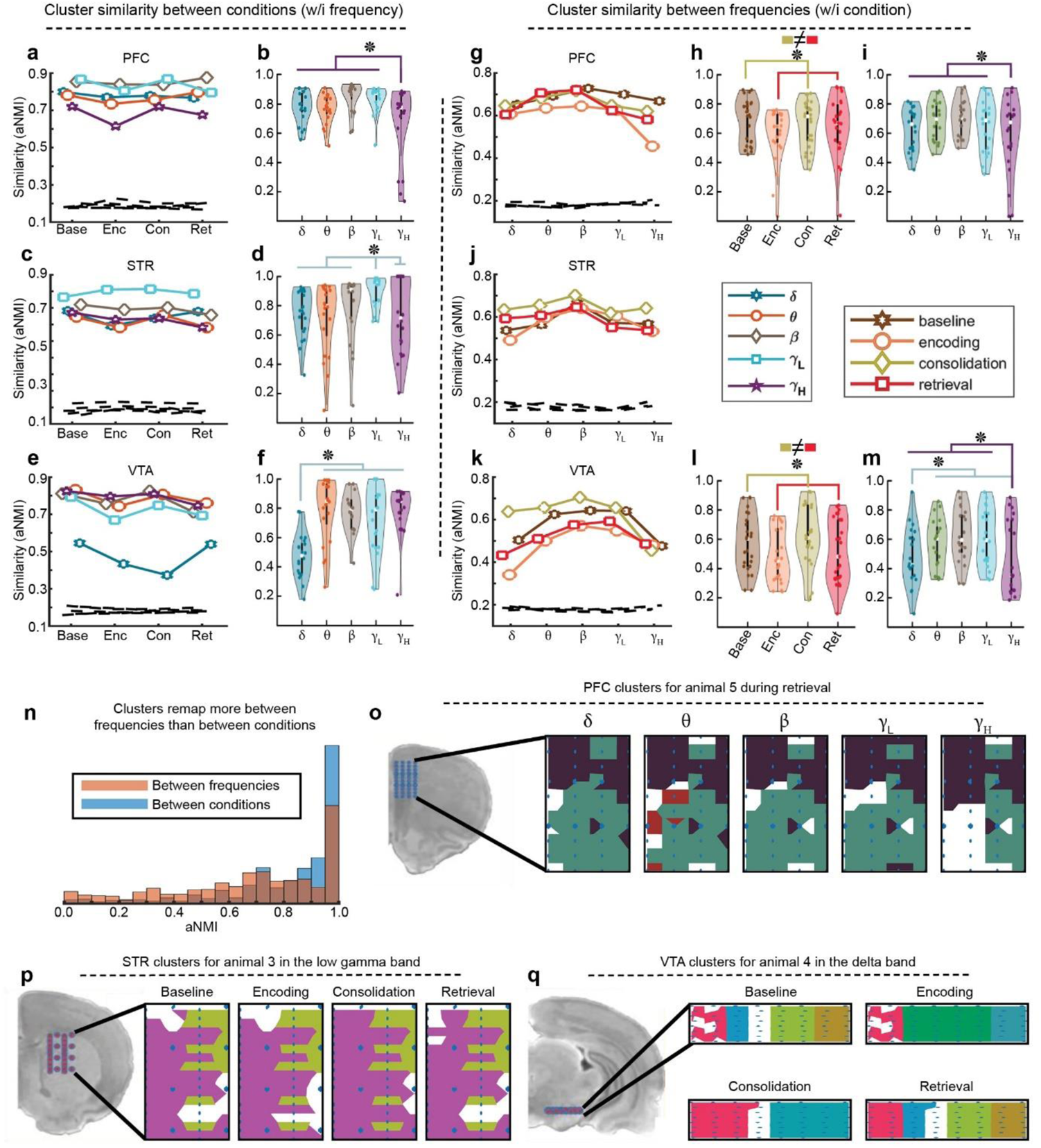
Intra-regional cluster stability. **a-f** Average normalized mutual information (aNMI) calculated across conditions but within frequency band. **a** PFC aNMI values (y-axis) are displayed for the four conditions (x-axis). Each line indicates results for a different frequency band (legend is next to panel **j**). Dashed lines indicate expected values in an analysis of random clusters. **b** PFC aNMI values were grouped by frequency band. Violin plots show aNMI values (y-axis) for condition similarity per frequency (x-axis). Each animal is represented by 4 dots (one for each condition) for each frequency. **c-d** Similar to **a-b** except for the STR. **e-f** Similar to **a-b** except for the VTA. Delta had lower aNMI than other frequency bands. **g-m** aNMI calculated across frequencies but within condition. **g** PFC aNMI values (y-axis) are displayed for the five frequency bands (x-axis). Each line indicates results for a different condition. **h** PFC aNMI was grouped by condition. Violin plots show aNMI values (y-axis) for frequency similarity calculated within each condition (x-axis). Each animal is represented by 5 dots (one for each frequency) for each condition. **i** PFC aNMI was grouped by frequency, generating a plot similar to **b**, except the underlying calculation here was within condition instead of within frequency. **j** Similar to panel **g** for data from the STR. **k-m** similar to **g-i** for data from the VTA. **n** Histogram shows aNMI in the between frequency (red) and between condition (blue) analyses. Between-frequency comparisons generally had lower aNMI. **o-q** Examples of cluster remapping. Anatomical location of recording array is shown to the left, and clusters are mapped to anatomical space in panels to the right. Pseudo colors indicate different clusters (unassigned channels have no color). **o** This example shows that high gamma in the PFC had a cluster scheme different from the other bands (see also lower aNMI values in panel **i**). **p** Clusters in the low gamma band in the STR. Note the stability in cluster organization across conditions (see also panel **d**). **q** This example demonstrates the effect observed in panel **e**. Namely, delta in the VTA had an unstable cluster map across conditions. Stars indicate significant linear contrasts in an ANOVA model.

In general, aNMI values were higher than would be expected by chance, but also consistently below 1, meaning that internal network structure in the PFC, STR, and VTA was neither completely remapped or completely stable either when looked at across different conditions (Figure 6a-f) or across different frequencies (Figure 6g-m). More specifically, for every paired combination of animal, condition, region, and frequency, we generated 1000 random pairs of cluster schemes where the total number of channels, the number of clusters, and the number of channels per cluster were held constant. aNMI between these pairs was calculated. The 99th percentile of these random distributions is plotted in Figure 6 (dashed lines). Random restructuring led to a maximum aNMI of about 0.2 across all situations. Yet, we observed aNMI values that were consistently higher than this.

To examine remapping between different conditions we calculated the aNMI of cluster schemes within frequency between different conditions. Separately for each condition, this analysis captures the average similarity of a condition with the other three conditions while holding frequency constant. For example, considering only clusters observed in the delta band in the VTA and averaging across animals, the NMI of the cluster organization observed during consolidation had similarities of 0.51, 0.25, and 0.35 with the clusters observed during baseline, encoding, and retrieval, respectively. Averaging these three values yielded 0.37 which is displayed in Figure 6e. aNMI values were submitted to a 5 (frequency bands) X 4 (conditions) within subjects ANOVA for each region (the ANOVA numerical data are presented in Extended data Figure 6-1; here we highlight only the relevant significant results). In the PFC there was a main effect of frequency (Figure 6a-b). Visual inspection of Figure 6a indicated that this effect was driven by reduced cross-condition stability in high gamma, and this was confirmed by linear contrast. But it appears that a small number of data points drove the effect (Figure 6b). In the STR there was also a main effect of frequency (Figure 6c-d), and this was driven by relatively high stability in the low gamma band (Figure 6d) as well as low stability in the theta band (not shown). The effects in the PFC and STR were relatively modest (η²<.2). By contrast, the VTA exhibited dramatic remapping of its cluster structure in the delta band (Figure 6e-f; η²=.48). Taken together, the STR was generally stable across conditions in all frequency bands. The PFC exhibited moderate restructuring of its cluster structure in the high gamma band, and the VTA restructured dramatically across behavioral states, but this restructuring was limited to delta band signaling.

To examine independence between different frequencies we again used aNMI. Separately for each frequency, this analysis captures the average similarity of cluster organization in one frequency band with the other four frequency bands while holding condition constant. These aNMI values were submitted to a 5 (frequency bands) X 4 (conditions) within subjects ANOVA for each region (for ANOVA table see Extended data Figure 6-2). In the PFC there were main effects of both frequency and condition (Figure 6g-i). These effects were driven by lower cross-frequency cluster scheme similarity in behavioral periods with an object present (both encoding and retrieval; Figure 6h) and lower cross frequency similarity in the cluster structure of high gamma signaling relative to other frequency bands (Figure 6i). There were no significant effects in the STR (Figure 6j). In the VTA there were main effects of both frequency and condition (Figure 6k-m). As in the PFC, behavioral periods with objects had lower cross-frequency cluster structure similarity than those without an object (Figure 6l). Also similar to the PFC, the cluster scheme for high gamma was dissimilar from the cluster schemes of other frequency bands. In addition, and unlike the PFC, the cluster structure of delta signaling was dissimilar to other frequency bands in the VTA (Figure 6m).

Taking these two analysis approaches together, PFC high gamma and VTA delta exhibited significant changes in cluster schemes. This indicates that these areas remap their internal structures with respect to signalling in these frequency bands across conditions (For single animal example see Figure 6q) and that the physical layout of signaling in these frequency bands is different from other frequency bands (for single animal example see Figure 6o). Furthermore, both regions exhibit more cross-frequency dissimilarity in the object periods, suggesting a greater degree of functional segregation between frequency-specific signal generators during object interaction. By contrast, the STR exhibited relatively high stability across conditions (for single animal example see Figure 6p). Finally, it is clear from visual examination of Figure 6 that cluster structures are generally more differentiated between different frequencies than across different conditions, suggesting independence of the neural substrates supporting signaling in different frequency bands. Averaging across animals, conditions, regions, and frequencies, within frequency aNMI values had a mean of .82 (Figure 6a-f), but within condition aNMI values had a mean of .68 (Figure 6g-m)(*p*<<.001; CI: .12-.16; see histogram in Figure 6n). That said, it should be emphasized that the most striking intra-regional cluster differences were observed within the delta frequency band in the VTA, suggesting that this structure remapped dramatically with respect to delta-band signal generation.

### Between-region network structure

As mentioned above, using GED to reduce the dimensionality of cluster signals to a single time course generally yielded a component that explained a sizable portion of the variance between channels (Figure 4f). After converting each cluster into a single time course, we examined connectivity between clusters using amplitude envelope correlations (Bruns et al., 2000). A bootstrapped null distribution was constructed for each connection (see methods).

Correlations were considered significant if they were stronger than 95% of their corresponding null correlations. The results of this procedure can be thought of as connectivity graphs for each animal in each condition. In these graphs, each node was a cluster with a specific region and frequency band, and edges were the correlations between nodes. Because there were often multiple clusters with the same region and frequency band, animals could sometimes have multiple connections along the same edge. In order to aggregate connections across animals, the strongest significant correlation between each frequency, region pair was taken for each animal. Any edge that did not have at least one significant connection for 4/5 rats was discarded. The medians of these maximum connection strengths across animals in each condition are plotted in Figure 7a-d. There were few significant connections including high gamma, so these connections were grouped with low gamma for this analysis. In general, the pattern of connectivity was dense in the baseline period, with 72% of all possible connections exhibiting significant coupling. Connectivity dropped during the encoding period, with 2% of all possible connections exhibiting significant strength. Connectivity then rebounded in the consolidation period to 52%, and then fell again during the retrieval period to 18%. In addition, while PFC delta was the node with the highest betweenness centrality in the first three behavioral conditions, PFC theta became the node with highest betweenness in the retrieval period (dot size in Figure 7a-d).

In order to unpack these results further, we replotted the connectivity as a function of change relative to connectivity strength during baseline (Figure 7f-h for increases; 7i-k for decreases). In general, connections in the theta band strengthened marginally during the consolidation and retrieval periods (Figure 7e; p’s<.07). In addition, there was also beta band connection strengthening in the consolidation period (p=.0503). In the retrieval period, a complex pattern of high frequency interactions involving the beta and gamma bands emerged (Figure 7h). Interestingly, decreases in specific delta band connections between regions were observed in all three conditions, but these were not significant in the aggregate (Figure 7e).

To check whether the aggregating process had biased the results, we performed an analysis of graph theoretic descriptive statistics on the full cluster X cluster connectivity matrices of significant connections derived for each rat. In general, nodes with high strength also had high betweenness, high clustering coefficients, and low path lengths (Figure 7l-o). We summed the strength of all clusters within each region and frequency band (Figure 7p-s). The results observed in the aggregated graphs were recapitulated. Overall strength reduced markedly in the object periods relative to the non-object periods. In addition, while strength was concentrated in the delta band during baseline (Figure 7p), theta band connections exhibited the most strength during the consolidation period (Figure 7r). These results were confirmed with a series of within-subject ANOVAs comparing pairs of conditions using frequency band, condition, and region as factors (for full ANOVA tables see Extended data Figures 7-1 to 7-3). Confirming the overall drop in strength during the encoding and retrieval periods, there was a main effect of condition in comparisons between baseline and encoding and between baseline and retrieval (Fs(1,281)>114; ps<<.001), but this main effect was absent when comparing baseline to consolidation (p=.8). Confirming the shift from delta to theta strength, there was an interaction between condition and frequency for comparisons between baseline and all three other conditions (Fs(4,281)>8.8; ps<<.001). Planned comparisons targeted at examining changes in relative delta/theta strength found that delta band connections were weaker in the consolidation period relative to baseline (t(58)=-3.3; p=.002), and connection strength in the theta band was marginally increased during the consolidation period relative to baseline (t(60)=1.9; p=.055).

We considered whether inter-regional connections played a role in memory. To test for this, connectivity strength of all significant connections was calculated for each session independently. Memory strength for each session was assessed as shown in Figure 1e. For each animal, we took the mean connection strength and memory strength values across sessions and then calculated the correlation between these values. Interestingly, during the consolidation period, theta connections between the VTA and STR and between the VTA and PFC were significantly correlated with memory (p’s<.05; Extended data Figure 7-1). However, a correlation analysis with only 5 animals should be interpreted with an appropriate amount of caution.

Finally, to check whether clustering had meaningfully contributed to our network connectivity findings at all, we repeated the GED and connectivity analysis considering entire regions as singular clusters (Extended data Figure 7-2). In general, this analysis found markedly fewer significant connections. In particular, only 3 connections were found in the retrieval period when entire regions were considered, compared to 26 connections observed using clusters. In other words, segregating the intra-regional activity into clusters was crucial to uncovering the memory-related functional dynamics.

## DISCUSSION

Rats spent less time exploring previously encountered compared to novel objects (Figure 1), and this memory effect was associated with a complex and dynamic pattern of inter-regional functional connectivity (Figure 7). At baseline, we observed that the STR, PFC, and VTA were robustly coherent across multiple frequency bands, with delta oscillations playing an outsized role. There was a dramatic pruning of network connectivity when rats were exposed to a novel object. After the novel object was removed, connectivity rebounded, but the connectivity profile shifted away from being dominated by delta towards being dominated by theta. Finally, when animals were re-exposed to objects, connections were not as severely reduced as they had been during initial presentation, and specifically theta and higher-frequency connections were stronger than they had been during the novel object encoding period. Underlying these inter-regional changes, functional organization of gamma frequency signals in the PFC and both gamma and delta signals in the VTA all changed markedly across behavioral conditions.

It is important to appreciate that these patterns were detectable only with the use of sub-regional clustering analysis (supplemental Figure 3). Although there was considerable variability in the signals recorded at different electrodes (Figure 2), we found that sub-regional clusters of electrodes were stable across multiple sessions recorded on different days. These clusters were verified using 20-fold validation, silhouette value examination (Figure 3), and by comparing the statistics of observed clusters to those of randomly chosen clusters (Figure 4). In general, clusters covered between a quarter and a third of the space of our electrode arrays (mean cluster size 18-24 electrodes; Figure 4g). Thus, for the STR and PFC, clusters covered an area of approximately half a square millimeter, and in the VTA they covered less than a tenth of a square millimeter. These areas are smaller than the traditional demarcations between architectonically categorized brain regions (Paxinos and Watson, 2006). This highlights the rich pattern of fine-grained spatiotemporal dynamics that can be discovered only through large-scale recordings and multivariate data analyses.

The idea that such small areas could act as functionally important units in long distance patterns of connectivity is consistent with principles of anatomy: Anatomical tract tracing studies have often found exquisite patterns such that regions lying only a single millimeter apart can have dramatically different profiles of connectivity (Schmahmann and Pandya, 2006), and the patterns of connectivity between our three recording targets are no exception (Prensa and Parent, 2001; Gabbott et al., 2005; Geisler and Zahm, 2005; Hoover and Vertes, 2007). Recent work has begun to reveal the functional importance of highly specific anatomy. For example, in rodents, specific fiber pathways are independently responsible for dopamine-dependent learning about novel objects and social stimuli in the VTA (Gunaydin et al., 2014). In monkeys, connectivity between small cortical patches supports face perception (Grimaldi et al., 2016; Chang and Tsao, 2017; Moeller et al., 2017). In humans, distinct subfields within the VTA are important for novelty and reward detection, and each of these subfields exhibits a unique pattern of functional connectivity (Krebs et al., 2011). The present results help to generalize these findings further, showing how sub-regional patches of brain tissue form changing patterns of long-distance connectivity during novel-object memory encoding, consolidation, and retrieval.

In addition, we observed that signals at different temporal frequencies and signals measured during different behavioral conditions both had distinct cluster topographies (Figure 6). This suggests that frequency-specific signal generators are anatomically localized and can be activated or deactivated depending on task demands, resulting in a constantly shifting landscape of functional anatomy. This finding also builds on earlier work. For example, in humans the BOLD activation associated with semantic concepts changes across the entire cortical mantle in response to attentional goals (Çukur et al., 2013), and nodes of the default mode network become less connected during cognitively engaging tasks (Raichle, 2015). More generally, Honey et al. (Honey et al., 2007) used a computational model of biologically inspired brain signals and known anatomical connectivity of the macaque brain to simulate electrophysiology data. They found that functional connectivity simulated over a long time window (minutes) recapitulated patterns of anatomical connectivity, but on shorter time scales (seconds or less) patterns of functional connectivity deviated from the model’s set anatomy. The authors interpreted this finding to mean that the brain is capable of dynamically changing its functional connectivity in ways that would not be predicted from anatomy alone, and our results confirm this interpretation. However, Honey et al. (Honey et al., 2007) reported that functional connectivity exhibits regression towards the mean over relatively short periods of time (10s of seconds). By contrast, we observed sustained periods with dramatically different cluster structures and long-distance connectivity, implying that both local and global network states can be held far from any equilibrium for at least several minutes in response to environment/task changes. As discussed in Honey et al. (Honey et al., 2007), computational modelling efforts with explicit consideration of context may capture this phenomenon.

Examining the specific pattern of connectivity changes exhibited in the present results, the lack of inter-regional connectivity during the encoding period is striking (Figure 7b and q). This result is surprising considering the vigorous novelty response produced by dopamine neurons of the VTA in both cats and monkeys (Ljungberg et al., 1992; Horvitz et al., 1997), and the finding that dopamine antagonists can impair memory in rodents (O’Carroll et al., 2006). In humans, dopaminergic single-unit firing in the substantia nigra has been shown to predict subsequent memory for novel stimuli (Kaminski et al., 2018). The seeming paradox of the known importance of DA in memory formation, juxtaposed with our observation of a disconnected VTA, could be explained by a connection between the VTA and an area that we did not record from. Much work has implicated the interaction between the HPC and VTA in response to novelty and memory encoding (Otmakhova et al., 2013). For example, fMRI data have revealed a novelty signal in the VTA associated with connectivity to the HPC, nucleus accumbens, and V1 (Krebs et al., 2011). The primary role for the HPC in the early stage of novelty encoding is further supported by faster neural response times for memory-predicting firing in the HPC compared with the substantia nigra in humans (Kaminski et al., 2018). Our data extend this finding by showing that other connections involving the VTA and important memory structures are suppressed during novelty encoding, heightening the importance of any HPC-VTA connection.

A second point of interest was the shift from delta (∼4 Hz) connectivity during the baseline period to theta (∼8 Hz) connectivity during the consolidation and retrieval periods (Figure 7e,p and r). There have been many reports highlighting coherent theta oscillations linking the HPC and PFC during declarative memory tasks (Benchenane et al., 2010; Otmakhova et al., 2013; Rajasethupathy et al., 2015; Kafkas and Montaldi, 2018; Kaminski et al., 2018), and putative DA cells in the midbrain of humans exhibit spiking coherence with PFC theta that is memory-dependent (Kaminski et al., 2018). By contrast, during a stimulus-response association task, delta frequency synchrony between the HPC, PFC, and VTA was interpreted as influence from the STR (Fujisawa and Buzsaki, 2011). This interpretation was based on prior observations of delta oscillations in the STR during this type of task. Where Fujisawa and Buzsaki (Fujisawa and Buzsaki, 2011) demonstrated that the HPC can be influenced by delta oscillations in a network involving the PFC and the VTA, our data demonstrate the reverse: the STR can be influenced by theta oscillations in a network involving the PFC and the VTA. We even observed some evidence that the theta connection between the VTA and the other two structures during consolidation was related to our behavioral measure of memory (Extended data Figure 7-1). Intriguingly, the largest changes in intra-regional cluster organization were observed for delta signaling in the VTA (Figure 6e). These changes may represent a state shift in the VTA from a delta-to a theta-influenced state. Indeed, it has recently been observed that theta and delta oscillatory modes in the HPC are orthogonal (Schultheiss et al., 2019), and our results indicate that these oscillatory modes may represent different network states beyond the HPC as well.

Finally, we observed a complex pattern of higher frequency connections during the retrieval period that were not present during the encoding period. It is widely accepted that memory retrieval involves a network of activation, and this is particularly true of old memories (Dede and Smith, 2018). Our data indicate that some network connections needed to support retrieval can be formed within minutes of initial encoding.

Three major limitations of this study are the need to relate the local clustering and global connectivity to single-unit firing, our lack of measurement of potentially involved structures beyond the STR, VTA, and PFC (primarily the hippocampus), and the need for more robust behavioral tests of memory. We believe these areas present important avenues for future work to extend the results presented here.

## Acknowledgments

MXC was funded by an ERC-StG 638589. NM and PMA were funded by a Hypatia award from the Radboud University Medical Center to MXC.

The authors declare no competing financial interests.

## Extended Data TABLES

**Extended Data Figure 6-1.**
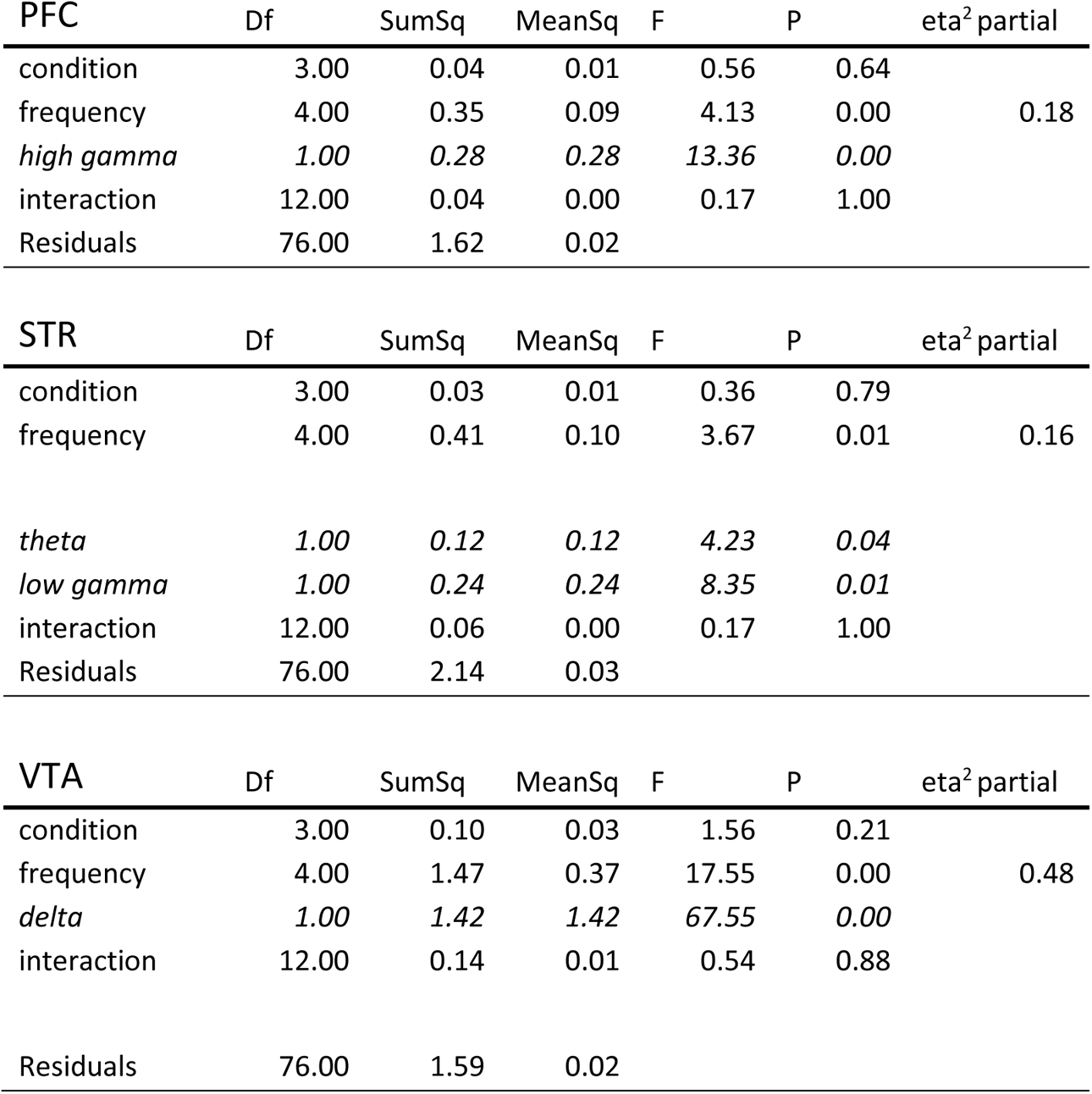
Cluster stability across conditions within frequency

**Extended Data Figure 6-2.**
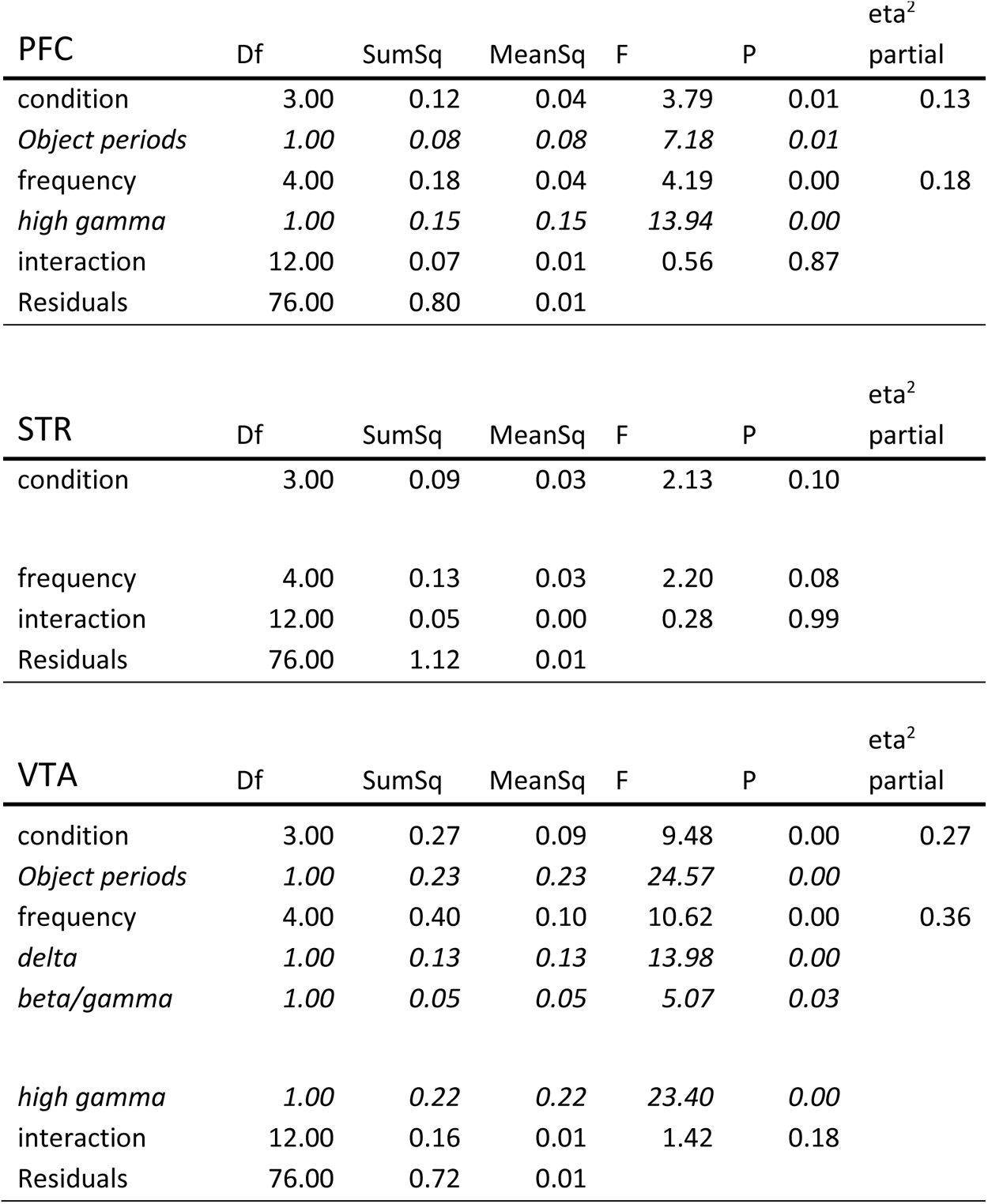
Cluster stability across frequency within condition

**Extended Data Figure 7-1.** Strength changes baseline vs. encoding

| Encoding | Df | SumSq | MeanSq | F | P | eta<sup>2</sup><br>partial |
| --- | --- | --- | --- | --- | --- | --- |
| cond | 1.00 | 4.52 | 4.52 | 114.26 | 0.00 | 0.29 |
| freq | 4.00 | 16.72 | 4.18 | 105.76 | 0.00 | 0.60 |
| reg | 2.00 | 0.30 | 0.15 | 3.82 | 0.02 | 0.03 |
| cond:freq | 4.00 | 2.05 | 0.51 | 12.96 | 0.00 | 0.16 |
| cond:reg | 2.00 | 0.16 | 0.08 | 1.99 | 0.14 |  |
| freq:reg | 7.00 | 0.85 | 0.12 | 3.08 | 0.00 | 0.07 |
| cond:freq:reg | 7.00 | 0.24 | 0.03 | 0.86 | 0.54 |  |
| Residuals | 288.00 | 13.04 | 0.05 |  |  |  |

**Extended Data Figure 7-2.** Strength changes baseline vs. consolidation

Baseline vs.
| Consolidation | Df | SumSq | MeanSq | F | P | eta<sup>2</sup><br>partial |
| --- | --- | --- | --- | --- | --- | --- |
| cond | 1.00 | 0.00 | 0.00 | 0.06 | 0.81 |  |
| freq | 4.00 | 25.56 | 6.39 | 141.14 | 0.00 | 0.66 |
| reg | 2.00 | 0.28 | 0.14 | 3.09 | 0.05 | 0.02 |
| cond:freq | 4.00 | 1.59 | 0.40 | 8.79 | 0.00 | 0.11 |
| cond:reg | 2.00 | 0.01 | 0.00 | 0.08 | 0.92 |  |
| freq:reg | 7.00 | 1.78 | 0.25 | 5.62 | 0.00 | 0.12 |
| cond:freq:reg | 7.00 | 0.52 | 0.07 | 1.63 | 0.13 |  |
| Residuals | 288.00 | 13.04 | 0.05 |  |  |  |

**Extended Data Figure 7-3.** Strength changes baseline vs. retrieval

| Retrieval | Df | SumSq | MeanSq | F | P | eta<sup>2</sup><br>partial |
| --- | --- | --- | --- | --- | --- | --- |
| cond | 1.00 | 4.22 | 4.22 | 121.23 | 0.00 | 0.30 |
| freq | 4.00 | 16.98 | 4.25 | 122.03 | 0.00 | 0.63 |
| reg | 2.00 | 0.11 | 0.05 | 1.54 | 0.22 |  |
| cond:freq | 4.00 | 2.10 | 0.53 | 15.10 | 0.00 | 0.18 |
| cond:reg | 2.00 | 0.08 | 0.04 | 1.13 | 0.32 |  |
| freq:reg | 7.00 | 1.05 | 0.15 | 4.31 | 0.00 | 0.10 |
| cond:freq:reg | 7.00 | 0.10 | 0.01 | 0.41 | 0.90 |  |
| Residuals | 283.00 | 9.84 | 0.04 |  |  |  |

---

**Extended data Fig 3-1.**
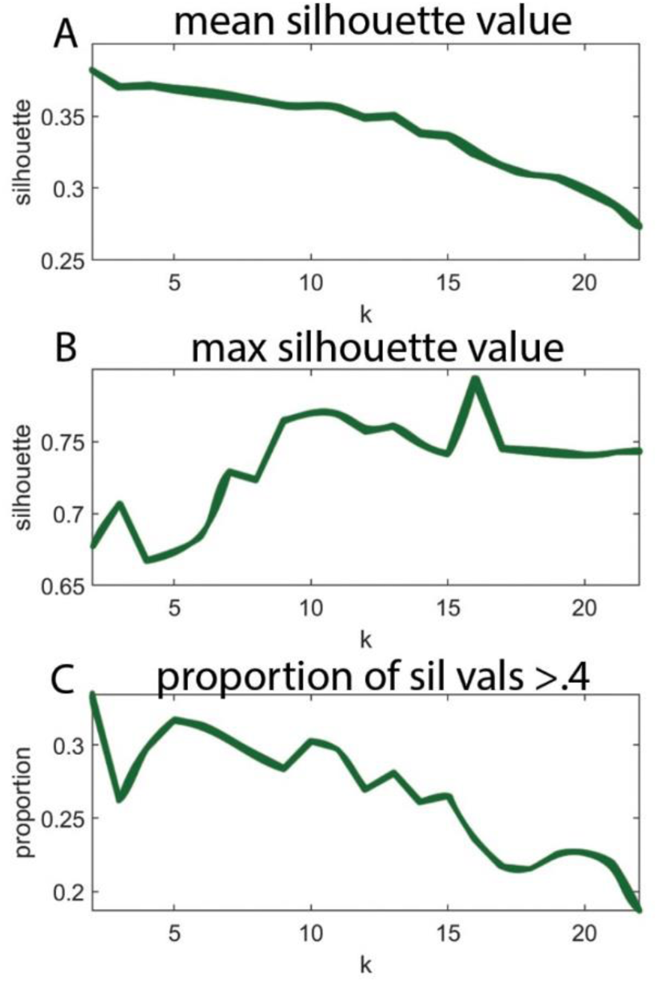
Choosing k for DBscan. **a** the mean silhouette value (y-axis) of clustering schemes calculated for all rats, regions, conditions, and frequencies using different values of k (x-axis). **b** The maximum silhouette value (y-axis) of clustering schemes calculated for different values of k (x-axis). **c** The proportion of silhouette values greater than .4 (y-axis) for different values of k (x-axis).

**Extended data Fig 7-1.**
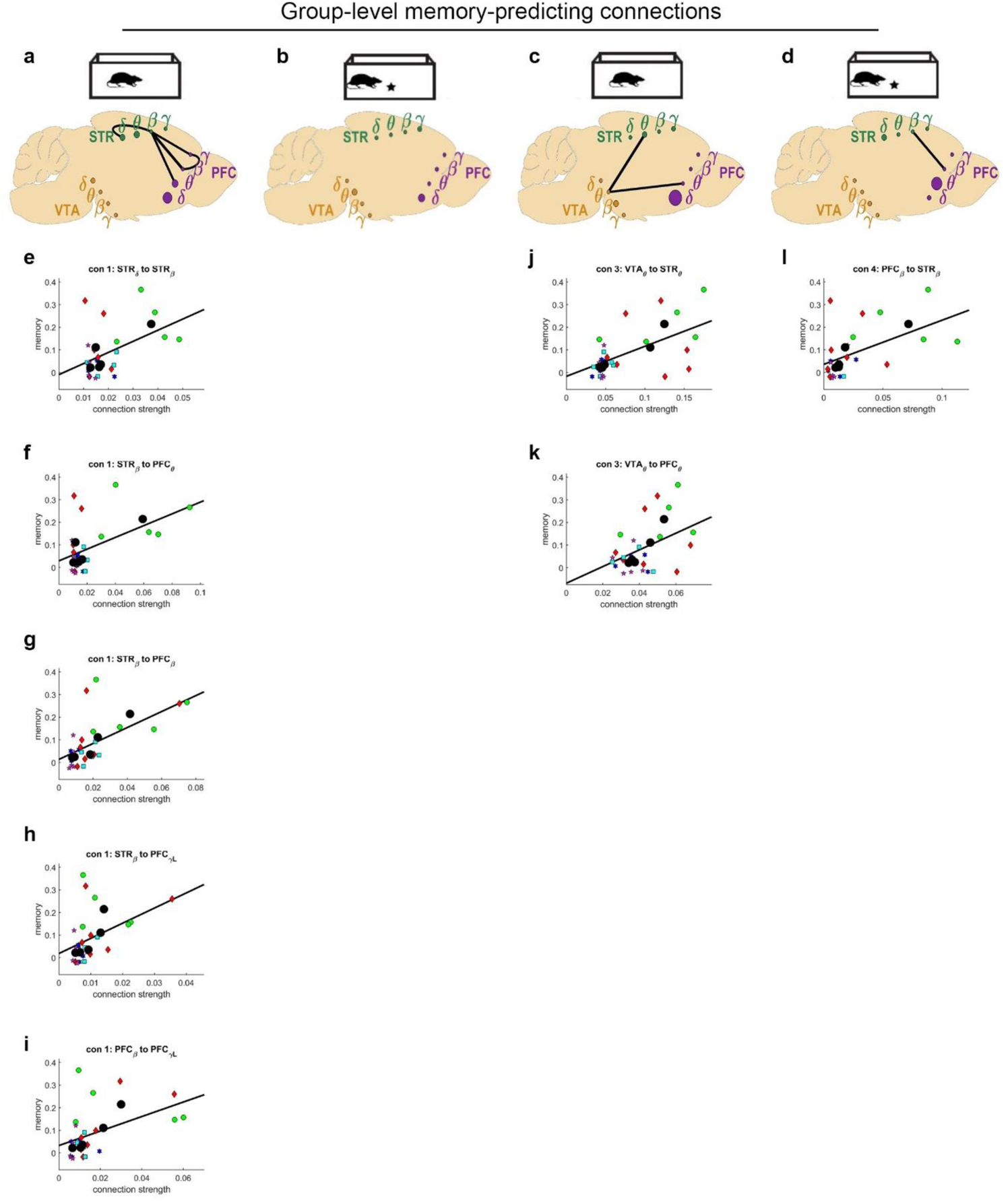
**Network connections related to memory.** For significant connections (main text Fig. 7a-d), we assessed the correlation between session connection strength and session memory. Memory was calculated as the proportion of time spent exploring the object during the encoding period minus the similar proportion for the retrieval period. There were two outlier sessions with memory <-.2 (main text Figure 1e), which were excluded from this analysis. **a-d** Network maps depicting connections that exhibited a significant correlation with memory strength. **e-l** Scatter plots depict the individual data points that went into all significant correlations. Before calculating correlations, each animal’s mean connection strength across sessions and mean memory strength across sessions were calculated. These animal mean values were submitted to correlation analysis. This analysis was also done using data from individual sessions in the correlation analysis. In general, a similar set of significant correlations were discovered. Different colors/shapes of points indicate the individual sessions for different animals. The large black circles indicate animal means. The regression line of best fit is shown (all ps<.05).

**Extended data Fig 7-2.**
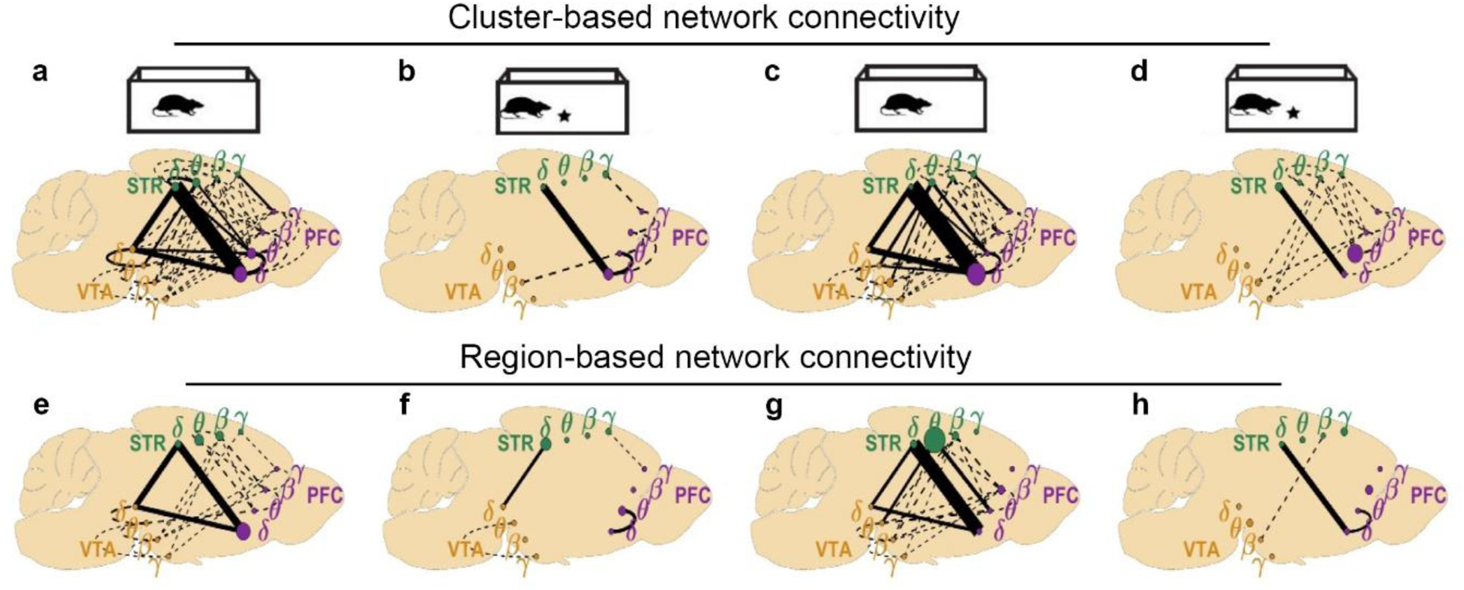
**Network connections between clusters versus between regions**. **a-d** Network maps are the same as panels **a-d** of Figure 7 (main text). **e-h** Network maps are calculated using all the same procedures as those in **a-d**, except each region was treated as a single cluster. Without considering the functional organization of signals within region (using clustering), many of the connections detected in panels **a-d** were missed in panels **e-h.** This is particularly evident during the retrieval period (panel h) where all of the complex high frequency interactions between regions have been missed.

